# Mechanism of proton-powered c-ring rotation in a mitochondrial ATP synthase

**DOI:** 10.1101/2023.08.11.551925

**Authors:** Florian E. C. Blanc, Gerhard Hummer

## Abstract

Proton-powered c-ring rotation in mitochondrial ATP synthase is crucial to convert the transmembrane protonmotive force into torque to drive the synthesis of ATP. Capitalizing on recent cryo-EM structures, we aim at a structural and energetic understanding of how functional directional rotation is achieved. We performed multi-microsecond atomistic simulations to determine the free energy profiles along the c-ring rotation angle before and after the arrival of a new proton. Our results reveal that rotation proceeds by dynamic sliding of the ring over the a-subunit surface, during which interactions with conserved polar residues stabilize distinct intermediates. Ordered water chains line up for a Grotthuss-type proton transfer in one of these intermediates. After proton transfer, a high barrier prevents backward rotation and an overall drop in free energy favors forward rotation, ensuring the directionality of c-ring rotation required for the thermodynamically disfavored ATP synthesis. The essential arginine of the a-subunit stabilizes the rotated configuration through a salt-bridge with the c-ring. Overall, we describe a complete mechanism for the rotation step of the ATP synthase rotor, thereby illuminating a process critical to all life at atomic resolution.

## Introduction

Adenosine triphosphate (ATP) is the energetic currency powering virtually all cellular processes. Eukaryotic cells produce most of their ATP by mitochondrial respiration. The oxidation of food stuff drives the pumping of protons across the inner mitochondrial membrane (IMM) into the intermembrane space, which results in the build-up of an electrochemical potential, the so-called proton-motive force. The ATP synthase enzyme complex (Figure 1a) sits in the IMM and harnesses the proton-motive force to catalyze the synthesis of ATP [1]. The present work is concerned with understanding how proton transfer across the IMM in turn drives the rotation of the transmembrane domain of ATP synthase.

**Figure 1:**
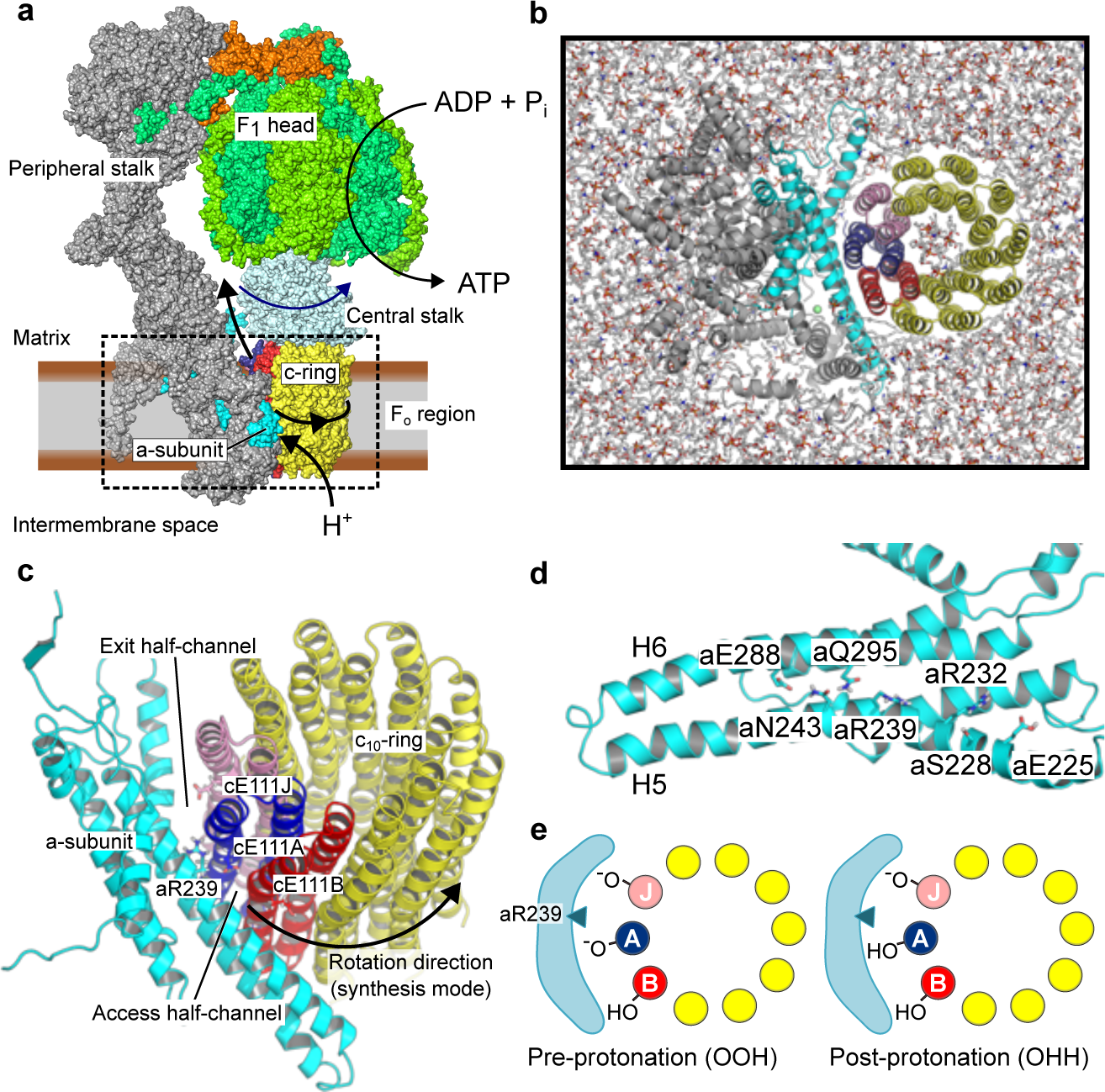
Overview of the system. (a) Structure and function of mitochondrial ATP synthase. The box encloses the F_o_ region investigated here. (b) Simulation model of *Polytomella sp.* mitochondrial ATP synthase F_o_ region, embedded in a realistic inner mitochondrial membrane (seen from F_1_). For clarity, solvent molecules and ions are not shown. (c) Side view of the a-subunit / c-ring complex highlighting important side chains at the interface. (d) Inner surface of the a-subunit highlighting the conserved polar residues. (e) Schematic of the two protonation states of the c-ring glutamates considered in this study. The drawings also illustrate the naming convention of c-ring subunits. Conventionally, the a-subunit is displayed in cyan; c-ring subunit J in pink; c-ring subunit A in blue; c-ring subunit B in red; all other c-ring subunits in yellow. This color scheme is used throughout the paper.

ATP synthase combines two molecular motors. The transmembrane F_o_ region channels the proton flow across the otherwise impermeable IMM in such a way that spontaneous proton-transfer powers unidirectional rotation of the turbine-shaped c-ring subdomain. The resulting torque is transmitted to the central stalk domain, whose rotation within the globular F_1_-ATPase domain triggers the conformational changes that power ATP synthesis. The peripheral stalk connects F_o_ to F_1_ and holds F_1_ in place. Given its central importance for cellular bioenergetics and the exquisite complexity of its two coupled rotary motors, ATP synthase has attracted considerable attention throughout decades of biophysical research [1, 2, 3]. Notably, the rotational mechanism of the central stalk and the F_1_ ATPase motor has been extensively studied [4], including by quantitative molecular simulations initiated from high-resolution crystal structures [5, 6, 7, 8, 9, 10].

More recently, the emergence of detailed structural information [11, 12, 13, 14, 15, 16, 17, 18, 19, 20, 21] has firmed up a general F_o_ rotational mechanism. The F_o_ rotor, or c-ring, rotates past the membrane-embedded a-subunit (Figure 1c). Their interface harbors two solvated half-channels for protons: the access channel on the intermembrane side and the exit channel on the matrix side. Crystal structures of isolated c-rings in lipid environment along with biochemical studies have shown that a conserved acidic residue (residue E111 of the c-ring subunit, or cE111 in short, in *Polytomella sp.*) is positioned to act as a proton shuttle. A proton binds to a negatively charged cE111 exposed in the access channel. Upon protonation, the now neutral glutamate can insert into the membrane, promoting rotation of the whole c-ring by one elementary step. Successive elementary rotations eventually lead the protonated glutamate to be exposed to the mitochondrial matrix in the exit channel. There, membrane voltage and higher pH conditions promote proton abstraction. Finally, one last elementary rotation puts the glutamate side chain back in the access channel.

Still lacking are a detailed and comprehensive picture of the substeps in the overall rotation mechanism and, even more importantly, a quantitative representation of the energy landscape underlying proton-driven c-ring rotation. Molecular Dynamics (MD) simulations of the c-ring at the atomistic and coarse-grained levels have advanced our understanding of the proton transfer process through the membrane and of the coupled rotation of the c-ring [22, 23, 24, 25]. However, structural information on the critical a-subunit has emerged only recently. Cryo-EM structures of full-length ATP synthase complexes at high-resolution, including the a-subunit, now open the way to a detailed analysis of the rotation mechanism by molecular simulations [26, 27, 28, 29, 30, 31, 32].

Here, we probe the structural rearrangements and free energy profiles along the rotation of the c-ring of *Polytomella sp.* mitchondrial ATP synthase in an all-atom F_o_ model embedded in a realistic IMM (Figure 1b) [33, 11, 16]. To understand how c-ring rotation is effected by the shifting protonation landscape of the a/c interface (Figure 1d), we compute the free energy profiles of an elementary c-ring rotation for the two possible protonation states of c-ring proton-carrying glutamates, namely before and after protonation (Figure 1e, Supplementary Table S1). Our results provide a plausible mechanistic and energetic description of the “electro-osmo-mechanical” step in ATP synthase operation.

## Results

Only two c-ring protonation states are likely to be relevant for proton-driven c-ring rotation (Figure 1e). The OOH state has the two adjacent c-ring subunits straddling the critical arginine unprotonated. By contrast, the OHH state has only one c-ring subunit unprotonated. By the 10-fold symmetry of the *Polytomella* c_10_-ring, the free energy surface of the OHH state is equivalent to the free energy surface of an HOH state shifted by 36 degree. The two relevant rotational free energy surfaces to be calculated, for the OOH and OHH states, are connected by proton transfer reactions, which will not be studied explicitly here. We assume that the rotary free energy surfaces are largely unaffected by any transmembrane potential difference because charges move parallel to the membrane plane in all processes modeled here.

### Free energy profiles along the c-ring rotation angle

We used geometric free energy calculations with the extended Adaptive Biasing Force (eABF) method [34, 35, 36, 37, 38] and a stratification strategy to estimate the potentials of mean force (PMFs) along the rotation angle *θ* of the c-ring (Figure 2), *i.e.,* the angle-dependent free energy profiles. To favor convergence of the calculations, we augmented the natural reaction coordinate *θ* with a second coordinate, namely the distance *d*_1_ between the cE111J CD atom and the aR239 CZ atom. *d*_1_ accounts for the possible formation of a salt-bridge between these two residues as rotation proceeds.

**Figure 2:**
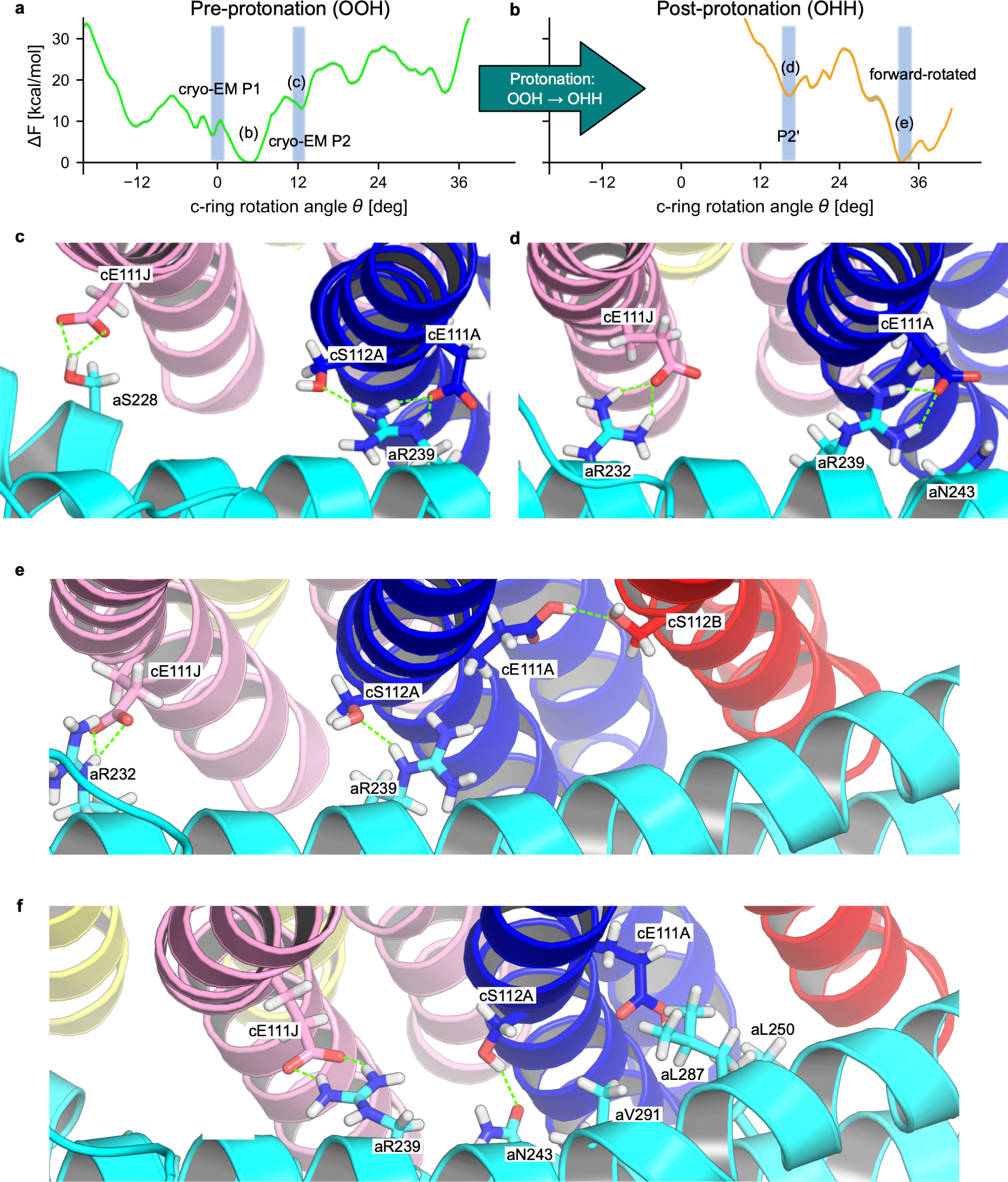
Free energy profiles and relevant configurations along c-ring rotation. (a,b). PMF along *θ* for the pre-protonation state OOH (a) and post-protonation state OHH (b). Grey outlines indicate ± statistical error estimated by bootstrap (see SI Appendix, Supplementary Figure S3). Shadings indicate relevant metastable states including cryo-EM structures P1 (PDB:6RD7/6RD9) and P2 (PDB:6RD8). (c) OOH ground state at *θ* ≈ 5^◦^. (d) OOH substate P2 at *θ* ≈ 12^◦^. (e) OHH substate P2’ at *θ* ≈ 16^◦^. (f) OHH forward rotated state at *θ* ≈ 36^◦^.

Our calculations, performed separately for the two relevant protonation states, OOH and OHH (Figure 1b-e), represent nearly ∼70 µs of all-atom MD simulation (Supplementary Table S2). The description and comparison of the profiles before and after protonation of the c-ring glutamate provide insight into the sequence of structural rearrangements, the plausible timing of proton transfer, and the energetics of the process.

For both relevant protonation states, the free energy profiles along *θ* exhibit several metastable states, including some that are separated by ≈36^◦^, *i.e.,* the size of an elementary rotary step (Figure 2a). Therefore, the PMFs are consistent with the expected spacing of the rotation free energy landscape, showing that our eABF calculations did pick up this important feature of the system. This point, along with the full, nearly uniform coverage of configurational space (Supplementary Figure S1c and d), the convergence of the force estimate (Supplementary Figure S2) and the low statistical error estimates (Supplementary Figure S3), supports the proper convergence of these challenging sampling problems.

#### Pre-protonation state OOH

State OOH exhibits a ground state at *θ* = 5^◦^, *i.e.,* with a slight forward rotation with respect to the reference configuration (Figure 2a). This finding indicates that already the deprotonation of state HOH contributes to the forward rotation of the c-ring. The reference cryo-EM structure P1, *i.e., θ* ≈ 0^◦^, corresponds to a metastable state and belongs to the same overall basin as the 5^◦^ state. From this basin, rotation in both directions entails an increase in free energy, corresponding to the cost of breaking the strong cE111A:aR239 interaction and inserting charged glutamate side-chains into the hydrophobic membrane. This is another expected feature of the free energy landscape which we correctly capture. Finally, a local minimum identified at *θ* ≈ 12^◦^ matches the so-called P2 alternate ring position observed in cryo-EM [33]. Overall, the free energy profile for our F_o_ system thus captures two key structures as local minima, indicating at least semi-quantitative consistency with experiments on the full F_o_F_1_ system.

To provide with a structural interpretation of the observed metastable states (Figure 2c and d), we computed the contact probability maps between key residues of the a and c subunits over the concatenated eABF trajectories. The contact maps of residues cE111 of c-ring chain A (thereafter, cE111A, which is followed by cE111B to cE111J around the ring) and cE111J with respect to polar a-subunit residues show that metastable states match reasonably well with specific, stabilizing contacts being formed (Supplementary Figure S4a,c,e). For example, the pre-protonation ground state at 5^◦^ is associated with a cE111J-aS228 hydrogen bond and a salt-bridge between cE111A-aR239, which are likely to be the main contributors to the stability of this substate (Figure 2c). Similarly, in typical configurations, substate P2 is stabilized by a cE111J:aR232 salt-bridge and interactions between cE111A, and aN243 and aR239 (Figure 2d), and is also compatible with configurations where cE111J replaces cE111A as the main interaction partner of aR239 (Supplementary Figure S1a). Therefore, the existence of a sequence of intermediates along the rotation pathway follows from the structural organization of the a/c interface, in which polar residues are positioned to sequentially form transient interactions.

#### Post-protonation state OHH

The overall minimum of the OHH rotational PMF is at *θ* ≈ 32^◦^ to 36^◦^ (Figure 2b). Considering the 10-fold symmetry, we can shift the OHH surface to the left by 36^◦^ to obtain the equivalent HOH surface, which thus has a minimum close to the most populated experimental angular state of 0^◦^. In addition, the post-protonation free energy profile exhibits several local minima. In comparison to the pre-protonation profile OOH, it shows a strong downhill trend to the forward-rotated state at *θ* ≈ 36^◦^ (Figure 2b). This downhill nature of the free energy profile results from the increasing electrostatic attraction between cE111J and aR239 (described by *d*_1_, see Supplementary Figure S1) as rotation proceeds, and is also consistent with the more favorable insertion of the now neutralized cE111A into the membrane. Thus, the difference between the OOH and OHH PMFs clearly shows how protonation effects rotation by shifting the free energy landscape, supporting our assumption that state OHH is primed for rotation.

The finer structure of the free energy profile, namely intermediates and barriers, nonetheless shows that the completion of the elementary rotation step does not proceed solely through relaxation along the free energy gradient, but involves a sequence of thermally activated transitions. Similar to the pre-protonation state, we used a contact map analysis to interpret these on-pathway intermediates in terms of a/c polar interactions (Figure 2e and f). We found that the final forward rotated state is stabilized by the cE:aR salt-bridge, established between the charged cGlu on the trailing c-ring subunit (cE111J) and the essential, absolutely conserved aArg239 residue of the a-subunit (2f). Additional stabilisation to the forward rotated state is provided by a hydrogen bond between aN243 and cS112A. Interestingly, in this state, the protonated cE111A side-chain still retains hydrating water molecules (Supplementary Figure S5c) and can either face towards the a-subunit inner surface or be in the so-called “ion-locked” configuration [39]. In this configuration, cE111A flips away from the a/c interface and forms a hydrogen bond with cS112B (Supplementary Figure S4d). A hydrophobic cluster on the a-subunit (aL250,aL287,aV291, see Figure 2f) may also contribute to the high free energy cost of rotating a charged c-ring subunit past *θ* ≈ 12^◦^ to 16^◦^.

Finally, we note that protonation destabilizes state P2, which is no longer a free energy minimum; instead, a novel minimum appears at *θ* =16^◦^, which we name P2’ (Figure 2e). Protonation of cE111A destabilises the cE111A:aR239 salt-bridge and makes cS112A the main interaction partner of aR239. As a result, the c-ring rotates forward by 4^◦^ and cE111A takes on the ion-locked configuration.

### Hydration of the a/c interface

In agreement with high-resolution cryo-EM structures [33, 20] and previous MD simulations [40, 28, 30, 31], we observe that water populates the access and exit half-channels all the way to the a/c interface (Supplementary Figure S5a). Water permeates the access half-channel through an opening between a-subunit H5 and H6 leading up to aH248 and aE288. Interestingly, the leading cGlu side chain (cE111A) remains solvated in the 36^◦^ forward-rotated state for both OOH and OHH (Supplementary Figure 5c and d). Furthermore, cGlu side chains buried deeply in the membrane are still marginally hydrated (C-H, see Supplementary Figure 5c and d). These observations suggest that desolvation of the protonated cGlu is not a strict requirement for membrane insertion as rotation proceeds. This may result in marginal transmembrane water flux, but not proton leakage because of the charge penalty. Lipid molecules, including cardiolipin, border the a-subunit on each side and may contribute to the interface (Supplementary Figure S6).

### Proton transfer and role of state P2

Whereas our classical simulations cannot directly describe the covalent chemistry of protonation of the c-ring acceptor residue cE111A, the analysis of interfacial water molecules in the eABF trajectories provides us with a plausible model for the timing of the proton transfer steps. The main result emerging from this analysis is that P2 is the most plausible rotational state for protonation of the c-ring to take place through a Grotthuss mechanism.

#### Direct proton transfer is unlikely

For *E.coli* ATP synthase, mutagenesis studies support aH245 as the proton donor [41]. In view of the conservation of the a-subunit [3], we assume an equivalent role for its *Polytomella sp.* homolog aE288 [33]. To assess the possibility of proton transfer by direct side-chain/side-chain interaction, we computed the 2-dimensional free energy profile *F* (*θ, d*_donor-acceptor_) (Figure 3a) by reweighting the eABF trajectories of state OOH (see SI Appendix, Supplementary Text). We also evaluated, as a function of *θ*, the (reweighted) average donor-acceptor distance. The results clearly demonstrate that direct contact between side chains aE288 and cE111 (*<*4 Å) is essentially never seen in the OOH state (Figure 3a). We conclude that the c-ring is not protonated directly as it rotates past aE288. As alternative, the proton may be transferred *via* water wires in a Grotthuss-type mechanism [42], as recently proposed by Spikes et al. [20].

**Figure 3:**
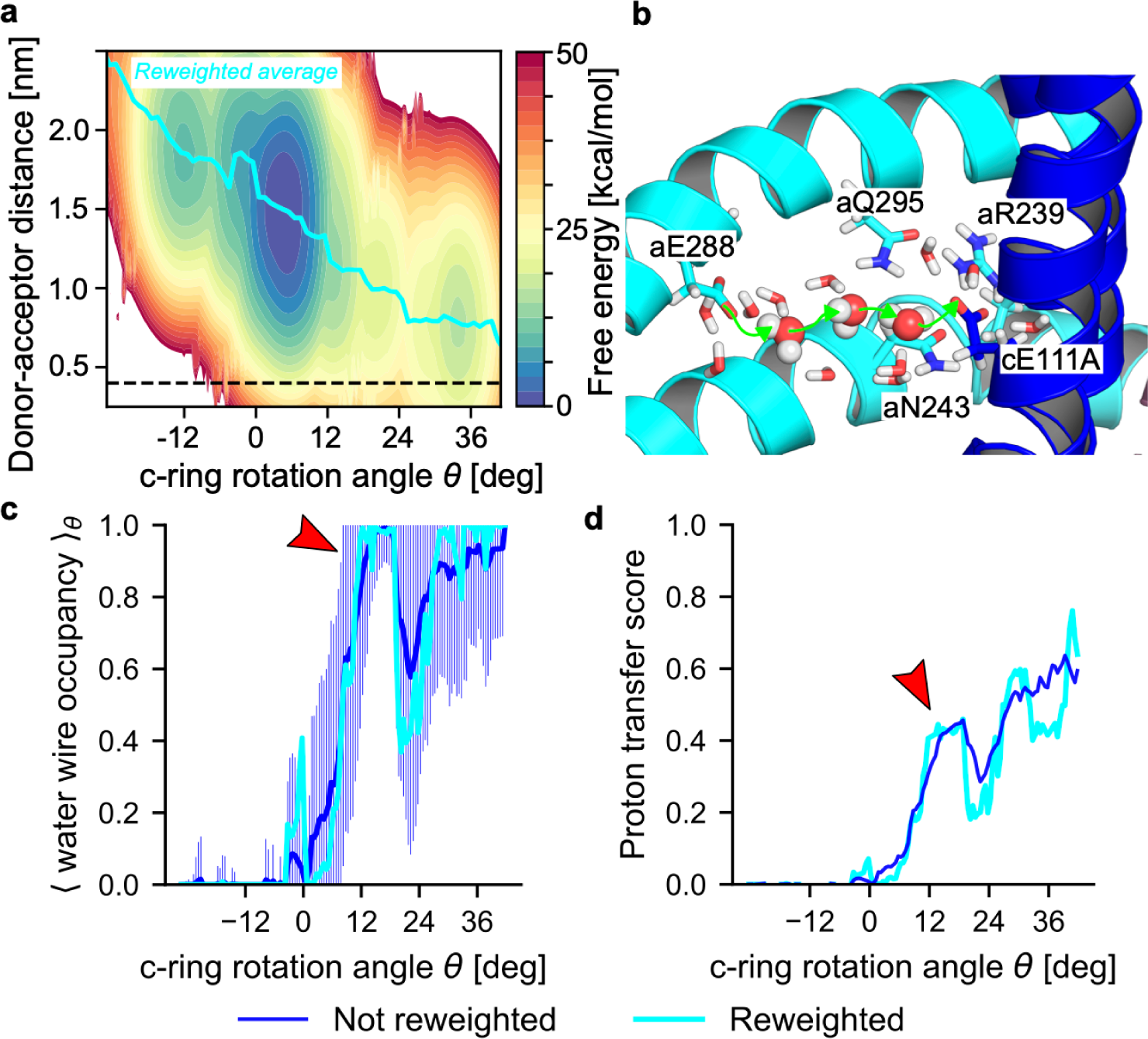
Facile protonation of the c-ring requires rotation into P2 state and water bridge. (a) PMF as function of rotation angle *θ* and distance between proton donor aE288 and acceptor cE111. The thick cyan line indicates the mean distance as function of *θ*, and the horizontal dashed line the distance where a direct proton transfer would be feasible. (b) Representative bridge of 3 water molecules between aE288 to cE111A in state P2. (c) Occupancy of water wires of length ≤ 8 in state OOH. Shown are the eABF-reweighted average (cyan) and non-reweighted average (blue) of the indicator function, which is equal to 1 if a wire of length ≤ 8 exists in the trajectory frame, 0 otherwise (shading: SD). (d) Water-mediated proton transfer score computed from eABF simulations of the pre-protonation state as function of *θ* (cyan: eABF-reweighted; blue: non-reweighted). Red arrows in (c,d) indicate state P2.

#### Grotthuss-type proton transfer along water wires is possible in state P2

Hydration of the acceptor residue (cE111A) is a necessary condition for water-mediated proton transfer, but is not sufficient. Water wires connecting the proton donor (aE288) and acceptor residues could provide a pathway for proton transfer. Therefore, we searched for water wires in the OOH trajectories that connect these two side chains by analyzing hydrogen bond networks between water molecules in their vicinity. This analysis clearly shows that water wires of length ≤ 8 are virtually always formed around state P2 (≈11^◦^ to 16^◦^), whereas they reach only ≈20 % occupancy in the ground state (Figure 3c). Further, wires of length 4 or lower are frequently observed for *θ* ≥ 12^◦^, whereas they are very rarely seen at lower values of the rotation angle (Supplementary Figure S5b). An example of a length-3 wire in state P2 is shown on Figure 3b. Although water wires are occasionally observed for negative *θ* states, they are very long (≥ 7 water molecules on average) and extend past the positively charged aR239 which separates the two half-channels, blocking proton leaks. Therefore, these wires will not be conducive to proton-transfer. Conversely, for *θ* ≥ 0^◦^, the leading c-ring subunit (A) is located in the access half-channel and proton transfer would thus not be impeded by aR239. In these conditions, proton-transfer along a single-file water wire is contingent upon every involved water molecule adopting a favorable orientation, which becomes rate-limiting, with the actual proton transfer occurring within femtoseconds [43]. Thus, the efficiency of proton transfer along a water wire is expected to decrease geometrically with wire length, provided a wire exists in the first place [44]. These considerations can be formalized into a proton-transfer score (see SI Appendix), which we used as a summarizing descriptor for the efficiency of c-ring protonation as a function of *θ*. The *θ*-dependent proton-transfer score evaluated from the eABF simulations of state OOH shows that P2 represents the first intermediate state to exhibit water wires both short enough to be conducive to proton transfer, and occurring with significant occupancy probability. We therefore conclude that water-mediated proton transfer is maximally probable in state P2, which ideally represents the proton-transferring state. Thus, we have an experimental structure for further examination of this important chemical step [33]. We also note that small but non-zero proton-transfer scores are observed for *θ* ≥ 5^◦^, suggesting a probabilistic picture in which proton transfer takes place over a range of angular states, with P2 nonetheless maximising the protonation probability.

### Dynamics of the Zn^2+^ cation

An electron density in the cryo-EM map of *Polytomella sp.* near residue aH248 was attributed to a Zn^2+^ ion, which takes slightly different positions in cryo-EM states P1 and P2 [33], see Supplementary Figure S7a. We performed blind prediction using a state-of-the-art convolutional neural network on the P1 structure, which recovered the assigned Zn binding site, supporting the attribution decision [45]. Because this cation may be involved in proton transfer [33], we investigated its positional dynamics and hydration shell in eABF simulations of state OOH (Supplementary Figure S7a-g). We found that the Zn^2+^ spatial distribution depends on *θ*. In the ground state, fluctuations of amplitude ≈5 Å in the *z* direction are observed (Supplementary Figure S7b), along with higher variance in the number of water molecules in the hydration shell (Supplementary Figure S7c). In P2, the fluctuations narrow down, and the cation interacts more closely with aE172 and aH248. Therefore, our findings suggest that the rotational state of the c-ring influences the positional dynamics of Zn^2+^, in qualitative agreement with the cryo-EM observation that the cation moves from P1 to P2.

## Discussion

We propose a complete mechanism of ATP synthase c-ring rotation, using all-atom free energy simulations representing nearly 70 µs of accrued simulation time. Our results shed light on important open questions about the proton-powered rotation of the F_o_ motor of *Polytomella sp.* mitochondrial ATP synthase. Because the general architecture of ATP synthase is conserved, we discuss our findings in light of recent structural, single-molecule and computational studies of various ATP synthase isoforms including *E.coli*, yeast mitochondria and *Bacillus sp.* PS3.

### Mechanism for proton-powered c-ring rotation

The rotation mechanism emerging from our analyses is summarised on Figure 4a. Initially, the motor is in the HOH state. In the exit half-channel, cE111J releases its proton, to bulk water or possibly a proton-accepting residue on the a-subunit such as aE225. The resulting OOH state rotates forward by *θ* ≈ 5^◦^ to 7^◦^ due to the rearrangement of polar contacts. The c-ring then undergoes angular fluctuations, until a thermally activated transition to state P2 is captured. This uphill transition over a significant free-energy barrier probably represents the rate-limiting step. We note that the barrier of 16 kcal mol^−1^ is likely overestimated because of imperfect sampling of the conformational dynamics. Then, protonation of the c-ring takes place in P2 along a structured water wire. This shifts the free energy landscape and destabilizes P2, promoting the rapid relaxation of the c-ring towards a metastable state at *θ* = 16^◦^. Finally, angular fluctuations allow the c-ring to cross smaller free energy barriers until the forward-rotated state is reached. Because of the large free energy gradient, this substep contributes most of the torque developed during c-ring rotation. Its driving force is the formation of the cE111J:aR239 salt-bridge, which provides a strong stabilization to the forward-rotated state, *i.e.,* HOH, and completes the elementary rotation step. Thus, this mechanism has features both of a powerstroke (down-gradient relaxation) and a Brownian ratchet (“capture” of forward rotational fluctuations) [46].

**Figure 4:**
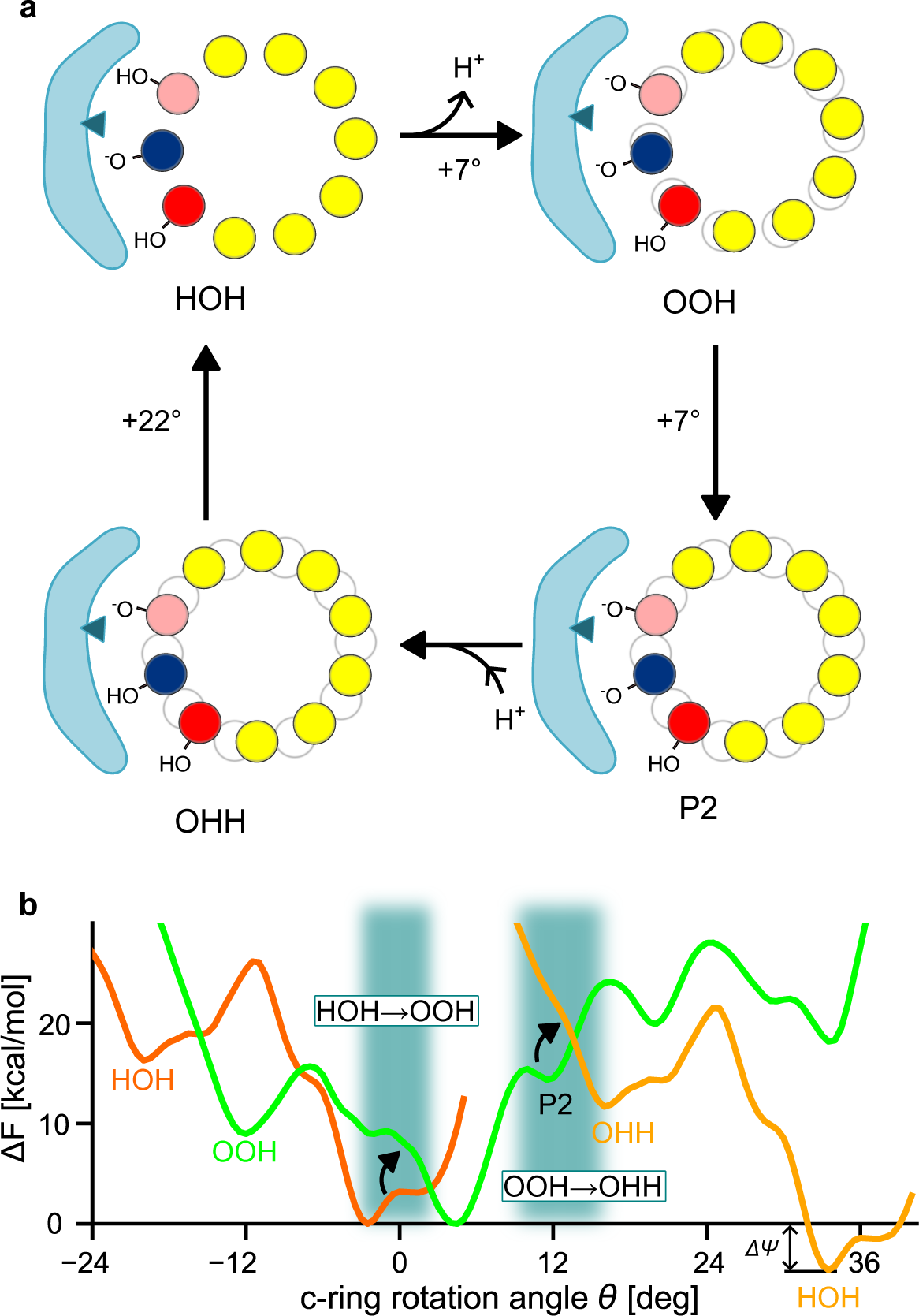
Mechanism of proton-driven F_o_ rotation. (a) Cycle of coupled proton transfer reactions and c-ring rotations. For easier visualisation of rotation movements, the grey outline materialises the initial position of the c-ring. (b) Free energy landscape of c-ring rotations coupled to proton transfer onto and off the rotating c-ring. Free energy profiles for states HOH and OHH are identical, up to a vertical shift by Δ*ψ* = 4.6 kcal mol^−1^ (*i.e.,* the value of the protonmotive force) and horizontal shift by 36^◦^. Shaded rectangles indicate proton transfer events.

Experimental mutagenesis identified aE288 as the final proton donor, suggesting that c-ring protonation may take place in a rotational state where cE111A and aE288 side chains are close enough for direct proton transfer [41]. However, our calculations suggest that such states may not be accessible from the pre-protonation state and instead support the growing consensus that incoming protons are transferred to the c-ring along a water wire through a Grotthuss mechanism [20, 40, 47]. Unlike previous computational studies of F_o_, which only evaluated global hydration of cGlu [28, 30], we specifically searched for water wires. Distinct short water wires between donor aE288 and acceptor cE111A indicate P2 as the likely state for proton transfer onto the c-ring. We note in passing that the proton-transfer score we introduce could be used to investigate water-mediated proton-transfer in other molecular systems. Upon transition to P2, the Zn^2+^ cation moves towards aE172, away from aH252 and is seemingly held more strongly. In P2, Zn^2+^ may provide an electrostatic barrier to prevent reverse transfer of protons to the bulk when the rotor is geared for protonation. Conversely, in P1, the increased positional freedom of Zn^2+^ may ensure that it does not hinder proton access to aH248 and aE288. The opening state of this “Zn-barrier” would be controlled by the competitive interactions with aE172 and aH252 (Supplementary Figure S7d-g). This hypothesis is consistent with Murphy et al.’s proposal that Zn helps synchronize proton access to the c-ring in the P2 proton-accepting state [33].

To keep the complexity manageable, the putative proton-donating (aE288, deprotonated) and proton-accepting (aE225, protonated) a-subunit residues were modelled in the same protonation state for both OOH and OHH, which were those predicted by our titration calculations but may not be representative of the true dynamics of proton transfers. Accounting for the charge changes upon protonation/deprotonation of these residues may alter the PMFs slightly. Nevertheless, the consistency of our model with independent experimental findings (see below) suggests that the protonation states of these residues do not majorly affect the details of the rotation mechanism [48]. Remarkably, when our protonation-dependent PMFs are shifted vertically by Δ*ψ* = 4.6 kcal mol^−1^ (corresponding to a 200 mV transmembrane electrochemical potential) and horizontally by ±36^◦^ (*i.e.,* the angular size of an elementary rotation), they intersect roughly at the presumed proton-transferring states including P2, suggesting that proton transfers are essentially iso-energetic (Figure 4b). Proton transfer would then move the system from the OOH to the OHH surface, where it first relaxes to a local minimum at about 16^◦^ before advancing to the new ground state near 36^◦^ (Figure 4b).

### Intermediates along the rotation pathway

Our results suggest that a/c interactions stabilise several intermediates along the rotation pathway, including the experimentally characterised P2. Thus, conserved, polar residues on the inner surface of the a-subunit (Figure 1d) are positioned to enable successive formation/breaking of interactions between a and c, thereby realizing a dynamic sliding of the c-ring onto the stator by stochastic jumps between metastable states. Whereas a large, negative free energy gradient would thermodynamically promote net directional rotation, it may not be sufficient to reach biologically useful timescales. Instead, by establishing smaller barriers between successive rotation substeps, on-pathway intermediates could facilitate rotation, as proposed for other biological and artificial molecular machines [49]. An additional role for the polar residues, which is also supported by our simulations, may be to help stabilise the water wires and thus assist with proton-transfer [18, 47], see Figure 3b.

### cE:aR salt-bridge

Among the conserved residues of the a-subunit, the critical arginine (aR239) is required for proper function of the proton-powered rotor [50, 51, 52, 53, 54]. The electrostatic interaction cE:aR between cE111 and aR239 was proposed to contribute to directionality [55]. A cE:aR salt-bridge is observed in some recent cryo-EM structures [20], but not all [17, 3], and its existence has also been challenged in a recent NMR study [56]. Our results show that a salt-bridge is formed nearly without interruption throughout rotation, either with the leading (A) or trailing (J) c-ring subunit. The transition between the two seems precisely to happen in state P2, which could help orient the cE111A side chain towards the water channel. In the OOH state, the competition between salt-bridges involving adjacent cE111 residues transiently stalls the rotor; protonation promotes rotation by favoring the formation of the aR239:cE111J interaction.

### Nature of the state primed for rotation

Whether a single-charged (OHH) or double-charged (OOH) ring-protonation state is primed for rotation is unclear. Triple-charged (OOO) intermediates were observed in coarse-grained simulations of c-ring rotation with dynamical protonation, lending credit to an OOO/OOH scenario [27]. Here, the comparison of the energetics of forward rotation reveals that only state OHH exhibits a large free energy drop for forward rotation, consistent with it being the primed state. By contrast, in state OOH, forward rotation clearly entails an increase in free energy, making forward-directed rotation from this state unlikely. Given the three charged cGlu, the PMF for state OOO would also exhibit steep climbs in free energy in both rotation directions. Therefore, in the OOO/OOH scenario, there is no state with the expected characteristics of the primed state. Our model relies on the hypothesis of OHH and OOH states being most relevant, which is supported by alchemical calculations on yeast ATP synthase indicating that OHH may be the dominant protonation state [28]. A recent solid-state NMR study found that up to 4 cGlu side chains are facing the a-subunit, suggesting that 4-times deprotonated (*i.e.,* OOOO) configurations are explored [57]. However, our results show that such “open” conformations are also compatible with protonated cGlu (see *e.g.* Figure 2e).

### Comparison with single-molecule studies

Recent single-molecule experiments on *E.coli* ATP synthase concluded that a 36^◦^ elementary rotation is broken up into a 11^◦^ protonation-dependent step, followed by a 25^◦^ rotation driven by electrostatic interaction [47, 58, 59]. Our results are remarkably consistent with these findings, whose mechanistic underpinnings they firm up. A minor difference is that we find the initial step to be split into two substeps, with deprotonation of the lagging c-ring subunit (HOH → OOH) contributing 5^◦^ to 7^◦^ and the protonation of the leading c-ring subunit (OOH → OHH) 5^◦^ to 7^◦^. The total extent of this step, 10^◦^ to 14^◦^, is consistent with the experimental measurement of 11 ± 3^◦^, all the more so that perfect agreement is not to be expected since the single-molecule experiments were performed with full-length F_o_F_1_ ATP synthase from *E. coli*. Both the sequence differences and the influence of the central stalk and F_1_ head may affect the details of the mechanism. Recent F_o_ high-resolution structures suggest that the *θ* = 10^◦^ to 14^◦^ substep is common in a range of species [33, 18], suggesting that this mechanism could be general.

### Comparison with earlier all-atom MD studies

Recently, free energy calculations along c-ring rotation were reported for yeast mitochondrial ATP synthase [28], and for *Bacillus PS3* ATP synthase [30, 31]. For yeast, state OOH was deduced to undergo rotational diffusion up to ≈10^◦^, which is in line with our findings. The interactions between aR (aR176 in yeast) and other a-subunit polar residues, and the c-ring glutamate (cE59 in yeast) were found to shape the rotational free energy landscape. The leading cGlu got progressively more hydrated as rotation proceeded from *θ* = 0, supporting an increase in protonation probability and an assignment of the ≈10^◦^ configuration as the probable proton-transferring state. Our findings agree with these conclusions. Yet, in contrast to our model, the authors propose that directional rotation stems from the free energy barrier to forward rotation being lower than that for backward rotation in the primed state (OHH) [28]. This proposal is based on umbrella sampling and could be consistent with recent theoretical analyses of molecular motors [60]. However, the yeast rotational PMF for state OHH shows a steep climb in free energy in both directions and no marked free energy minimum at the forward rotated state. To achieve a decreasing free energy in the synthesis direction, these investigators had to deprotonate c-ring subunit I; however, this site is buried in the membrane and thus unlikely to be accessible for deprotonation. The resulting state is equivalent to our OOH state rotated by −36^◦^. Although this state would indeed rotate forward, this movement would not result in a cycle of rotations that can be closed. Thus, this state is unlikely to be representative of a functional rotational state, leaving the rotation mechanism unclear.

1 µs-long MD simulations of *Bacillus* PS3 F_o_ led to the conclusion that the Coulombic attraction between the deprotonated trailing cGlu and aR in state OHH is the main driving force for c-ring rotation [30]. Further, these investigators deduced that the HOH → OOH →OHH transitions occur with minimal angular change [30]. The subsequently reported free energy profile [31] for state OHH exhibits a negative gradient from −20^◦^ to 0^◦^, compatible with forward rotation. However, no on-pathway intermediate was detected (but note that the reference c-ring *Bacillus* PS3 structure [19] (PDB: 6N2D) is actually rotated by ≈10^◦^ with respect to the *Polytomella* reference (PDB: 6RD7/6RD9), see Supplementary Figure S8). Here, by using the aR239:cE111J salt-bridge distance as an auxiliary reaction coordinate for free energy calculations, we arrive at free energy profiles both for OHH and OOH states that are consistent with forward rotation in a closed cycle (Figure 4) and capture a key intermediate consistent with structural [33, 20] and single-molecule studies [61, 58, 47, 59].

### Importance of lipids

We observe that lipids, including one cardiolipin, populate the outer a/c interface on both the IMS and matrix sides (Supplementary Figure S6), possibly contributing to strengthen a/c interaction as recently proposed for *E. coli* [21]. Previous investigators have suggested that cardiolipin molecules stably bound in the vicinity of the a/c interface may play a role for functional rotation [24, 62]. Our observations are consistent with this proposal, but our sampling timescales may not allow for the complete equilibration of the lipid distribution around the *F_o_* domain. Therefore, the presence of cardiolipin near the a/c interface may reflect in part the initial arrangement of the membrane rather than increased affinity of F_o_ for cardiolipin. It has also been proposed that the central lumen of the c-ring in F- and V-ATPase is populated by lipids [63, 64], but their nature and stoechiometry is unknown in *Polytomella*. Whereas previous investigators modelled POPE [65, 28] or POPC [30, 31] in the lumen, we used cardiolipins, which may also contribute to explaining the different rotational PMFs. Future studies will have to address the interplay between lipids and the energetics of c-ring rotation.

## Conclusion

The functional, directional rotation of the c-ring is the starting point for the synthesis of ATP by ATP synthase. It is the result of a complex interplay between protonation/deprotonation, local dynamics of interactions, and large-scale subunit motion. We used atomistic free energy calculations to describe the structural mechanism and energetics of an elementary rotation step of the c-ring. Future studies will focus on integrating this structural description with a thermodynamically and kinetically consistent description of proton transfer to achieve a synthetic description of “osmo-mechanical” transduction by the F_o_ rotor.

### Data Archival

Simulation parameter files and all-atom structural models for representative configurations shown in Fig. 2 are deposited in the Zenodo repository 10.5281/zenodo.8124466 and will be made accessible upon publication.

## Materials and methods

### Preparation of structural models for MD simulation

Models for states OOH and OHH were prepared from the *Polytomella sp.* ATP synthase cryo-EM structure (6RD9) by keeping the c-ring, a-subunit and part of the peripheral stalk. Protonation states were determined by a multisite titration approach. Each model was embedded in a realistic IMM including cardiolipin, solvated in an orthorhombic box of TIP3P water molecules with 150 mM NaCl, and energy-minimised under harmonic restraints. Then, NVT equilibration (*T* = 300 K) was run for 10 ns followed by NPT equilibration (*T* = 300 K*, P* = 1 bar) for 20 ns while harmonic restraints were progressively relaxed, except on the peripheral stalk. Details of the equilibration procedure and simulation parameters are given in SI Appendix.

### MD simulations

MD simulations were run with GROMACS 2020.4 [66] using the CHARMM36m force-field [67]. See SI Appendix for detailed parameters.

### Extended ABF free energy calculations

eABF calculations were run from the coordinates and velocities of the equilibrated structures using the *colvars* module [68] with GROMACS. Absolute harmonic restraints were applied on the CA atoms of the peripheral stalk; therefore, to avoid absolute-restraint-related artefacts, we ran these calculations in the *NV T* ensemble (*T* = 300 K). One eABF calculation was run per model (*i.e.,* one for OOH and one for OHH). Two-dimensional PMFs were computed along *θ* (*i.e.,* the rotation angle of the c-ring with respect to the cryo-EM structure, with structural alignment on the a-subunit) and *d*_1_ (*i.e.,* the distance between cE111JCD and aR239CZ). To promote convergence of the calculations, an exploratory run was followed by stratified sampling [69]. eABF simulations amount to a total of 68 µs. One-dimensional PMFs along *θ* were obtained by Boltzmann-integration of the two-dimensional PMFs. Reweighting of eABF simulations was done as detailed in SI, using SciPy [70, 71] for interpolation and scikit-learn [72] for Kernel Density Estimation. See SI Appendix for details of the eABF protocol, convergence, and error analysis.

### Water-mediated proton transfer analysis

Water wires connecting residues aE288 and cE111A were identified by analysing all stratified OOH simulations with the breadth-first algorithm implemented in MDAnalysis [73, 74]. When applicable, the shortest water wire for a given frame was then identified using Djikstra’s algorithm as implemented in networkx [75]. The proton transfer score was defined as *k*(*θ*) = *k_0_ρ*(*θ*)*α*^⟨^*^n∗^*^⟩^*^θ^* with *k*_0_ = 1, *ρ*(*θ*) the occupancy (*i.e.,* existence probability) of at least one water wire of length ≤ 8 at c-ring rotational angle *θ*, ⟨*n*^∗^⟩*_θ_* the average length of the shortest water wire at c-ring rotational angle *θ* (using a conditional average restricted to frames were at least one water wire is present) and *α* = 0.8 an attenuation parameter, chosen to obtain appreciable signal. We derive the score and discuss the choice of *α* in SI Appendix.

### Data visualisation and rendering

Molecular structures were visualised and rendered with VMD [76] and Pymol [77]. Graphics were rendered with Matplotlib [78] and Inkscape [79].

## Acknowledgments

This work was supported by the Max Planck Society. We thank Bonnie Murphy and Werner Kühlbrandt for stimulating discussions and the Max Planck Computing and Data Facility for providing computational resources. F.E.C.B. thanks Jérôme Hénin for advice about eABF, and Adrien Cerdan for critical reading of the manuscript.

## SI Appendix

### Detailed Methods

#### Atomistic models of ATP synthase F_o_ region in a realistic Inner Mitochondrial Membrane

##### Protein model

To study the rotation of the c-ring, we built a minimal model of the F_o_ region by extracting the c-ring, the a-subunit and part of the peripheral stalk from the recently published 2.8 Å-resolution cryo-EM structure of *Polytomella sp.* ATP synthase [33] (PDB:6RD9), see Main Text Figure 1b and Table 1. The missing loop of the a-subunit (residues 205 to 219) was reconstructed with MODELLER [80] using available lower-resolution coordinates as a template [16]. The missing C-terminal residues of the a-subunit were built *de novo* with MODELLER. The electronic density near aHis248 was modelled as a single Zn^2+^ ion.

**Table 1:** F_o_ region minimal model.

| Subunit | Chain | Residues |
| --- | --- | --- |
| c<sub>10</sub> | A to J | 54 to 127 |
| a | M | 95 to 327 |
| ASA1 | 1 | 520 to 617 |
| ASA3 | 3 | 154 to 321 |
| ASA5 | 5 | 1 to 61 |
| ASA6 | 6 | 28 to 151 |
| ASA8 | 8 | 2 to 89 |
| ASA9 | 9 | 1 to 97 |
| ASA10 | 0 | 2 to 82 |

**Table 2:** List and length of eABF simulations.

| Protonation state | Exploratory run | Stratified run | Total simulation time |
| --- | --- | --- | --- |
| OOH | 8.2 μs | $15 \times 1.64 = 24.6 \mu$s | 32.8 μs |
| OHH | 6 μs | $12 \times 2.48 = 29.7 \mu$s | 35.7 μs |

**Table 3:** Lipid composition of the membrane.

| Lipid | Proportion |
| --- | --- |
| Cardiolipin | 16 % |
| POPE | 37 % |
| POPC | 41 % |
| POPI | 6 % |

##### Protonation states

Protonation states were assigned according to the following protocol. First, residues cE111A and cE111J were assigned the standard, negatively charged state (*i.e.,* the system was modelled in state OOH, see Table 4). Then, the most probable protonation states as a function of pH for all other titratable residues (ASP, GLU, LYS, HIS) were determined by multisite titration p*K_A_* calculations [81]. The protein interior, solvent and membrane were treated as continua of dielectric constants 4.0, 80, and 4.0 respectively. The membrane was modelled as a 3.5 nm-thick slab. Ions in the solvent were modelled by a Boltzmann-distributed charge density corresponding to 150 mM NaCl at 300 K. APBS [82] with the tAPBS front-end was used to solve the Poisson-Boltzmann equation for the electrostatic potential. Then, protonation probabilities as a function of pH were evaluated by Monte Carlo sampling with the Karlsberg2 program [83, 84]. When appropriate, solvent-exposed residues were assigned the protonation state corresponding to the pH of their membrane compartment, *i.e., pH* = 6.8 for the IMS and *pH* = 7.7 for the matrix. Notably, this resulted in residues aE288 and aE225 being protonated. For all titratable residues on c-ring subunits (except cE111) a consensus protonation state was determined by the majority rule. To model state OHH, we added a proton on cE111A, but did not perform a new p*K_A_* calculation, that is, we neglect the possible impact of changes in protonation of cE111A on other titratable residues. This ensures that the protonation state of cE111A is the only changing parameter between models. Structural models were prepared for MD simulation using CHARMM.

**Table 4:** Protonation states of the c-ring.

| State | Label | Protonation states |
| --- | --- | --- |
| Pre-protonation | OOH | cE111J: charged, cE111A: charged, cE111B: protonated |
| Post-protonation | OHH | cE111J: charged, cE111A: protonated, cE111B: protonated |

##### Membrane model

A membrane patch with average composition of cauliflower inner mitochondrial membrane [85] (see Table S1) was built using CHARMM-GUI [86, 87, 88] and equilibrated by 20 ns of molecular dynamics with GROMACS. Cardiolipin (CDL) molecules were modelled in the CDL^2–^ state. The protein was inserted in the membrane by deleting overlapping lipids. Cardiolipin molecules in the central hole of the c-ring were kept; clashes between them and the c-ring atoms were relieved by energy-minimizing them in a cylindrical harmonic potential centered on the c-ring axis (using the MMFP module of CHARMM).

##### Preparation for molecular dynamics simulation

The system was embedded in an orthorhombic box of dimensions 15.1 nm × 12.5 nm × 17.5 nm and solvated with TIP3P water molecules. Na^+^ and Cl^−^ ions were added to ensure electroneutrality of the system and achieve an approximate salt concentration of 150 mM. GROMACS-format topology files were obtained using the psf2itp.py program from the CHARMM-GUI suite [89]. Each system was energy-minimized with absolute harmonic restraints on the protein CA atoms, the Zn^2+^ ion, and lipid head-groups (force constant 4184 kJ*/*mol*/*nm^2^).

#### Molecular dynamics simulations

##### General parameters

The energetics was modelled with the CHARMM36m force-field [67] with a van-der-Waals cutoff of 1.2 nm and using van-der-Waals force-switching from 1 nm. Electrostatics were treated by the Particle Mesh Ewald method for the long-range component, and the cutoff between short-range and long-range was set to 1.2 nm.

All molecular dynamics simulations were performed with GROMACS 2020.4 [66] using the leapfrog integrator with a 2 fs timestep. Covalent bonds involving hydrogens were constrained using LINCS. In all production simulations, the peripheral stalk residues were restrained close to their initial positions using absolute harmonic restraints on CA-atoms (force constant 4184 kJ*/*mol*/*nm^2^). Temperature was maintained at 300 K using the velocity-rescale thermostat [90] with a coupling time of 100 ps on three separate temperature groups (protein atoms, membrane atoms, water and ions). During *NPT* equilibration, pressure was maintained at 1 bar using a semi-isotropic Berendsen barostat (*xy* and *z*) with a time constant of 1 ps and compressibility 4.5 × 10^−5^ bar^−1^ in both directions.

##### Equilibration protocol

After energy minimisation, each model was equilibrated according to the following 4-step protocol designed to progressively relax the membrane and protein.

1. 2 ns of *NV T* equilibration (*T* = 300 K, Berendsen thermostat) with harmonic restraints on the protein backbone heavy atoms and the heavy atoms of all lipid head groups.
2. 4 ns of *NV T* equilibration (*T* = 300 K, Berendsen thermostat) with harmonic restraints on the protein backbone heavy atoms, the Zn^2+^ ion and the heavy atoms of the *z*-coordinate of all lipid head groups.
3. 4 ns of *NV T* equilibration (*T* = 300 K, Berendsen thermostat) with harmonic restraints on the protein backbone heavy atoms and the Zn^2+^ ion, but all lipid restraints removed.
4. 20 ns of *NPT* equilibration (*T* = 300 K, Berendsen thermostat, *P* = 1 bar, Berendsen barostat) with harmonic restraints on the peripheral stalk backbone heavy atoms.

##### Extended Adaptive Biasing Force (eABF) calculations

###### Simulation setu

eABF calculations were performed to explore functional c-ring rotation. The use of an adaptive biasing force (ABF) enhances the sampling along two Collective Variables (CVs) by applying a biasing force. This biasing force is adaptively updated to compensate for the generalized thermodynamic force felt by the CVs. Thus, at convergence biased CVs undergo purely diffusive dynamics. At any time, numerical integration of the collected biasing force enables the estimation of the free energy landscape. Here, 2D integration was performed by Poisson integration as described in reference [38]. In eABF, ABF dynamics is applied on a virtual degree of freedom harmonically coupled to the CV [37]. Two-dimensional eABF calculations were run in *NV T* conditions starting from the temperature- and pressure-equilibrated systems, using the colvars module [68] interfaced with Gromacs 2020.4 and with active positional restraints. The *NV T* ensemble was chosen to avoid artefacts due to the usage of absolute positional restraint. To limit non-equilibrium effects, the *fullSamples* parameter was set to 10 000; that is, the bias is not applied at a given point of configurational space until this point has been visited at least 10 000 times to ensure a robust initial estimate of the bias. eABF was applied on *θ*, *i.e.,* the c-ring rotation angle with respect to its initial position, and on *d*_1_*J* (called *d*_1_ in Main Text), *i.e.,* the separation distance between cE111J CD and aR239 CZ. *θ* was defined using an orientation quaternion (spinAngle collective variable) computed on the CA atoms of residues (55 58 61 64 67 70 73 76 79 82 85 88 91 95 98 101 104 107 110 113 116 119 122 125) for each c-ring subunit, and using the *Z*-axis (0, 0, 1) as rotation axis [68]. The calculations were performed on a rectangular grid, with *θ* between −47^◦^ and 41^◦^ in 0.5^◦^ increments, and *d*_1_*J* ∈ 25 nm to 2.5 nm in 0.025 nm increments. In this way, we covered two full 36^◦^ rotation steps, one in the forward direction and the other in the reverse direction. Both CVs were harmonically coupled to extended degrees of freedom undergoing Langevin dynamics at temperature *T* =300 K, using force constants of 10 kcal*/*mol*/*deg^2^ for *θ* and 10 kcal*/*mol*/*Å^2^ for *d_1_J*. The Corrected *z*-averaged restraint (CZAR) estimator was used to correct free energy gradient estimates [37]. To keep the c-ring aligned with the *Z*-axis, we applied harmonic restraints on two virtual atoms, respectively close to the c-ring central axis’ top and bottom. Specifically, the centers of geometry of the CA atoms of residues (90 96) for all c-ring subunits, and of CA atoms of residue 59 for all c-ring subunits, were harmonically restrained (force constant 10 kcal*/*mol*/*Å^2^) to their respective position in the equilibrated structure. This mimics the presence of the central stalk in full-length ATP synthase. The eABF bias was occasionally observed to induce irreversible local unfolding of *α*-helices. When this happened, the simulation was backtracked until slightly before unfolding and relaunched with harmonic walls on the backbone inter-atomic distances. The harmonic wall potential *U_wall_*(*d*) applied on a backbone O N distance *d* is given in equation S1.

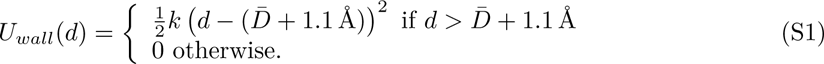

In equation S1, *k* = 20 kcal*/*mol*/*Å^2^ and 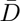 is the average distance during the first 50 ns of eABF simulation. Harmonic walls preserve local helix flexibility while preventing dramatic, irreversible unfolding. To promote convergence of eABF sampling, we used the two-step strategy introduced in reference [69]. After an initial long exploratory eABF run, CV-space was divided into non-overlapping windows and an eABF simulation was launched in each window starting from configurations sampled in the exploratory run, with the eABF bias accumulated in the exploratory run. Specifics of domain definition (Supplementary Figure S1c and d) and sampling times (Supplementary Table 2) differ slightly between states OOH and OHH.

###### Convergence and error analyses

Convergence of the eABF calculations was evaluated by monitoring the coverage of CV-space (Supplementary Figure S1c and d) and the Root Mean Square Deviation (RMSD) of the eABF force estimate (*i.e.,* the estimate of the free energy gradient) (Supplementary Figure SS2). The RMSD was computed separately for the uncorrected and CZAR-corrected gradient estimates. Most windows exhibit a stable gradient-RMSD in the last ≈100 ns of stratified eABF simulation, indicating that the bias has reasonably converged. The statistical error was estimated using a bootstrapping procedure based on the standard error of the mean of the accumulated biasing force, see Supplementary Figure S3 [91, 69]. We note that this procedure yields the error associated to the uncorrected potential of mean force rather than the CZAR-corrected ones. In absence of a rigorous procedure to propagate statistical errors through the CZAR-estimator, we chose to use the uncorrected errors to quantify uncertainty on the CZAR-corrected profiles. Estimated statistical errors do not exceed 0.2 kcal mol^−1^, indicating a low variance of the force-estimate which, together with the coverage of the relevant regions of configurational space (Supplementary Figure S1c and d), suggests proper convergence of the eABF calculation.

##### Reweighting of eABF calculations

Estimating the free energy landscape along degrees of freedom other than *θ* and *d*_1_*J* requires reweighting to correct for the eABF bias. Reweighting of eABF calculations is challenging because of the non-stationary bias. To our knowledge, there exists no rigorous procedure for this purpose. Nevertheless, approximate reweighting of orthogonal degrees of freedom can be achieved using the final, converged PMF to compute unbiasing factors - similar to an approach previously used for metadynamics [92]. With *A*_∞_(*θ, d*_1_*J*) the converged free energy profile from ABF, the applied biasing potential is −*A*_∞_. Thus, the (approximately) unbiased weight of configuration *x_i_* from the eABF trajectory is:

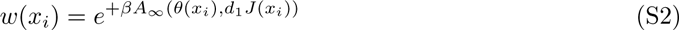

where *β* = 1*/k_B_T*. If configuration *x_i_* has unbiased probability *P* (*x_i_*) (*e.g.* in a long, unbiased equilibrium trajectory) and probability 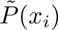 in the (converged) ABF simulation, then:

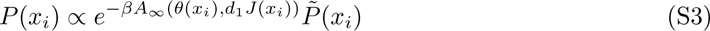

We implemented this reweighting procedure as follows. First, a bivariate spline is fitted to the PMF estimate, which makes it possible to compute the reweighting factor for any point in (*θ, d*_1_*J*) space. Spline fitting is performed using the RectBivariateSpline function of SciPy [70, 71]. Second, this spline approximation is used to compute reweighting factors (equation S2) which are then used in a weighted Kernel Density Estimation function, as implemented in scikit-learn [72]. For the reweighted PMF shown in Main Text Figure 3a, we used Gaussian kernels of bandwidth 0.1 nm for the donor-acceptor distance, and 1^◦^ for the c-ring rotation angle *θ*. Reweighting factors can also be used in weighted average calculations to unbias canonical averages, as is done in Main Text.

#### Water bridge analysis

Water bridge analysis was performed using the dedicated module in MDAnalysis version 2.0.0 [73, 74]. In brief, this module uses a breadth-first algorithm to iteratively search for water-mediated hydrogen bond bridges between selected residues. The length of shortest water bridge was computed with Djikstra’s algorithm in networkx [75].

#### Derivation of the proton transfer score

We define an effective proton transfer score to evaluate the proton transfer efficiency as a function of the rotational state of the c-ring. Inspired by reference [44], we reasoned that the proton transfer requires either a direct hydrogen-bond connection or the existence of at least one bridging water wire, and that proton transfer efficiency attenuates geometrically with the number of water molecules in the wire. For a given donor/acceptor/water configuration *x_i_* (*i.e.,* the Cartesian coordinates of the system, *e.g.* a simulation frame), this corresponds to the proton transfer efficiency model given in equation S4.

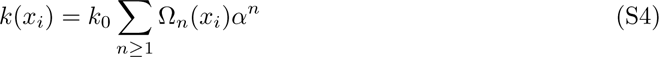

In equation S4, *k*_0_ is a basal efficiency, Ω*_n_*(*x_i_*) is the number of distinct water wires of length *n* at configuration *x_i_*, and *α <* 1 is a dimensionless attenuation factor quantifying the decrease in proton transmission probability as a new water molecule is added to the wire. Note that direct proton transfer could also be taken into account by extending the sum to *n* = 0 and setting Ω_0_ = 1 if direct proton transfer is possible, 0 otherwise. Here, we focus on water-mediated proton transfer and thus exclude the *n* = 0 term from the sum. Because of the geometric attenuation, the sum in equation S4 will be dominated by *n*^∗^(*x_i_*), *i.e.,* the smallest *n* for which Ω*_n_*(*x_i_*) ̸= 0. We thus use the approximation

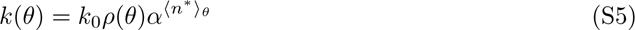

where *ρ*(*θ*) ≡ ⟨I*_n_*_≥1_(*x_i_*)⟩*_θ_* is the water-wire density with I*_n_*_≥1_(*x_i_*) is an indicator function taking value 1 if at least one water wire of length *n* ≥ 1 exists in configuration *x_i_*. The conditional averages ⟨· · · ⟩*_θ_* are over all frames *x_i_* with given values of *θ*. The values of ⟨*n*^∗^(*x_i_*)⟩*_θ_* were determined from eABF simulations in state OOH (with or without eABF-reweighting), as shown in Supplementary Figure S5b.

This formulation allows us to use directly the observables *ρ*(*θ*) and ⟨*n*^∗^⟩*_θ_* from eABF simulations, taking *k*_0_ = 1 since we are not concerned with the precise value of the score. Main Text Figure 3d shows the score computed from eABF simulations of state OOH (with or without eABF-reweighting) using *α* = 0.8, chosen to obtain appreciable signal. Supplementary Figure S5c illustrates the influence of *α* on the score profile. *α* is expected to depend in complex fashion on the translational and re-orientational dynamics of water molecules within a wire [44, 43].

Since our study focuses on comparing relative proton transfer efficiency to identify proton-transferring states, the approximations are appropriate, because they are not expected to dramatically change the qualitative picture. In the future, usage of the full expression for the score (*i.e.,* an averaged version of equation S4) and a careful parametrisation of *k*_0_ and *α*, for example from *ab initio* Molecular Dynamics simulations, could enable quantitative investigations of proton transfers in ATP synthase and other systems.

---

**Figure S1:**
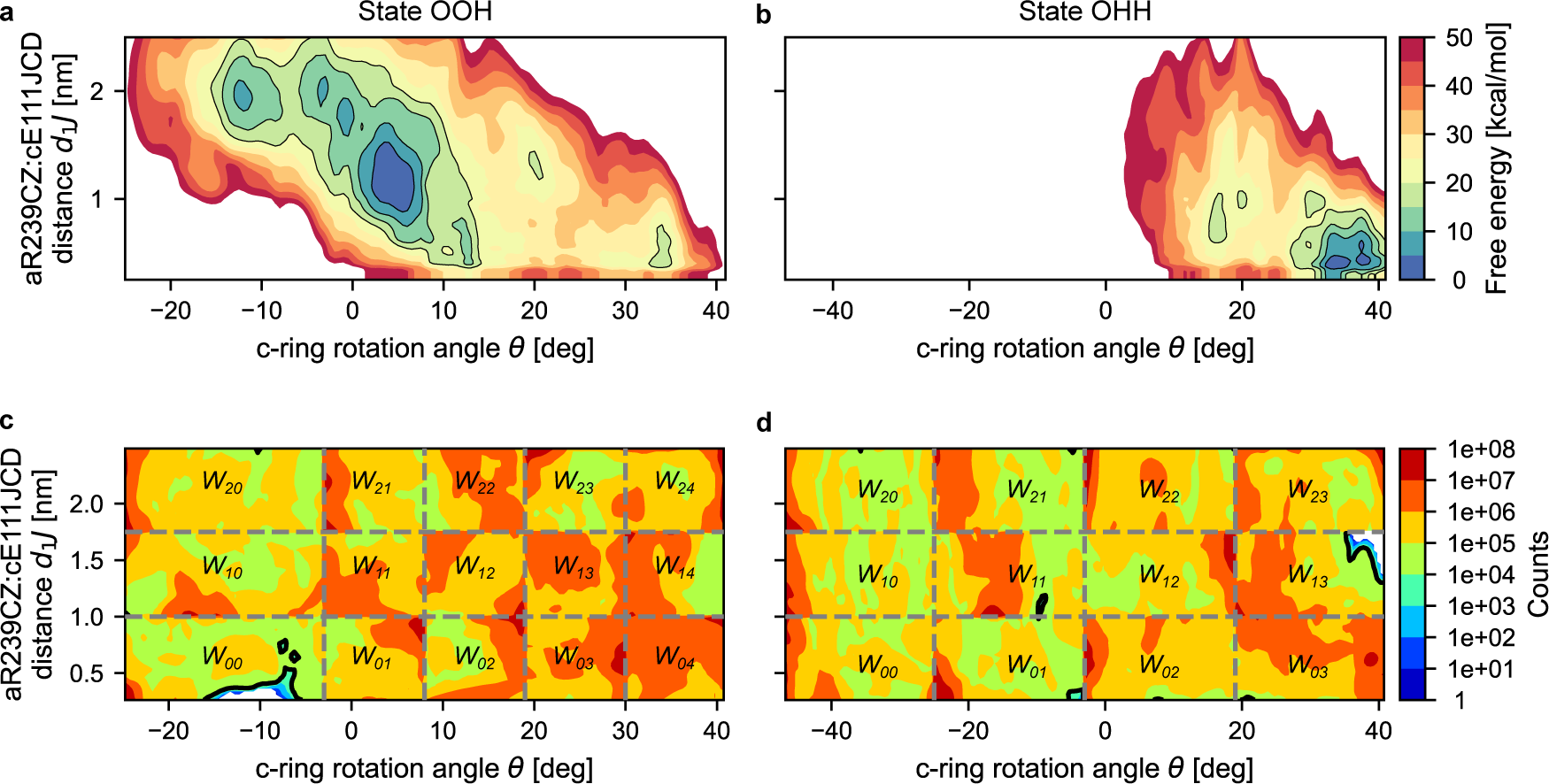
2D free energy landscapes computed by eABF. (a) Pre-protonation state (OOH). (b) Post-protonation state (OHH). (c-d). Sampling of the configurational space by eABF simulations. Shown are the respective counts in the 2D bins.

**Figure S2:**
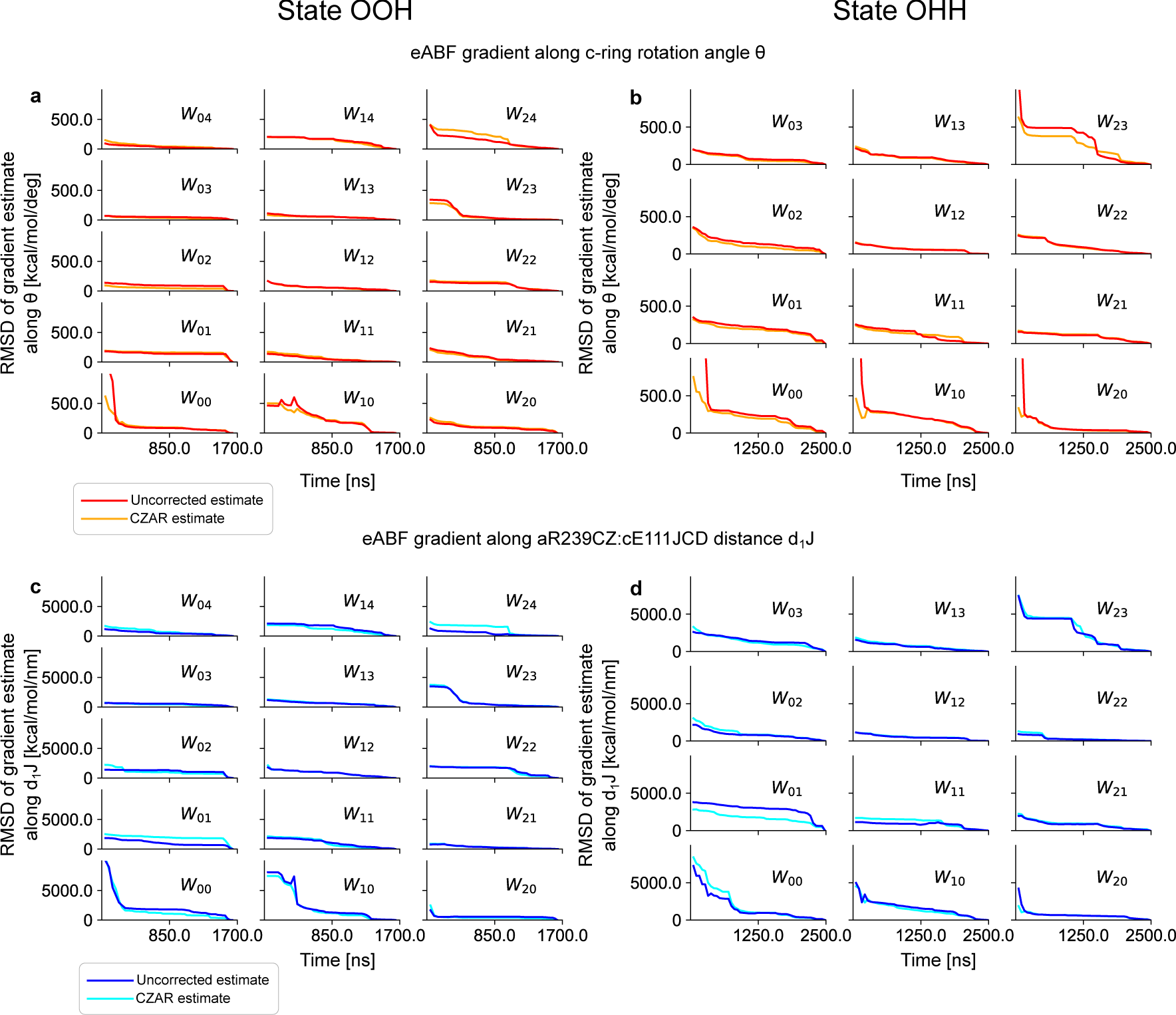
Convergence of eABF force estimate in stratified eABF simulations. In most windows, the gradient-RMSDs have stabilised in the last ≈100 ns, indicating that the eABF bias is reasonably converged.

**Figure S3:**
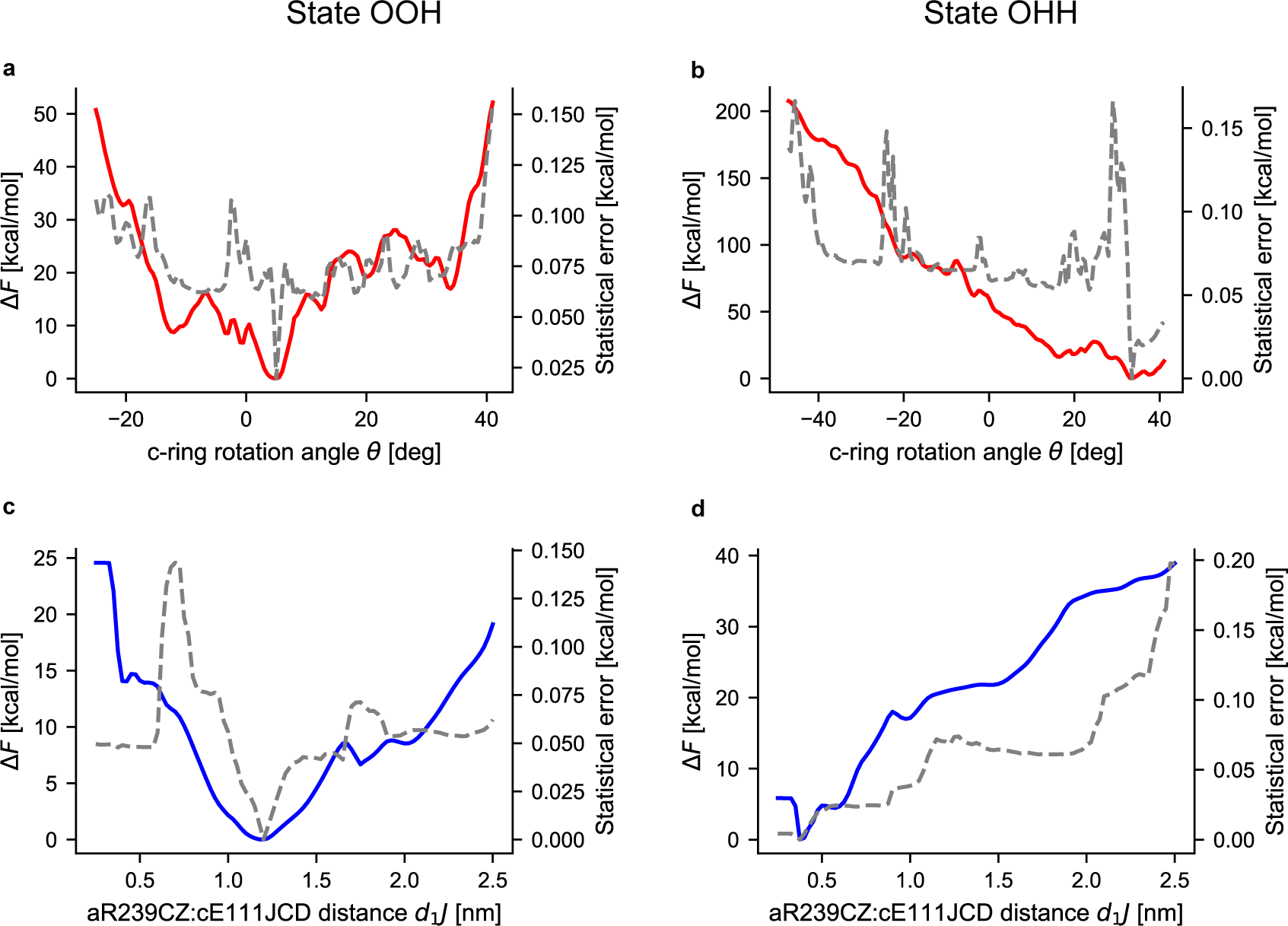
Bootstrap error analysis of eABF simulations. In panels (a) to (d), the free energies (left scale) are shown as solid colored lines, and the statistical errors of the PMF estimated by bootstrap are shown as dotted grey lines.

**Figure S4:**
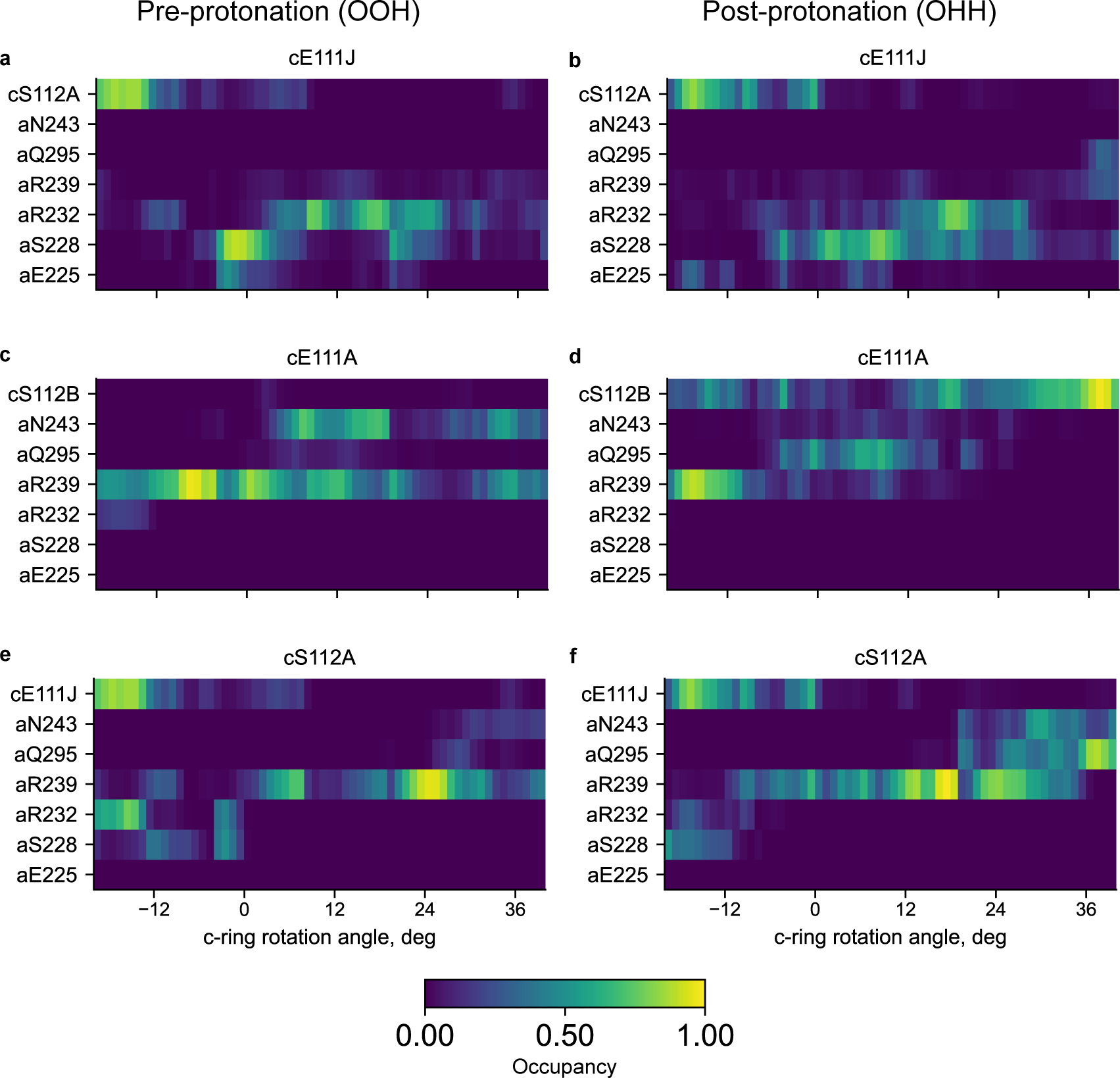
Contact map analysis of the polar interactions between c-ring and a-subunit in eABF simulations. The color indicates the probability of a particular contact to be formed at a given c-ring rotation angle (scale at bottom). Note that the weak signals observed along the aR239 row in panels a and b come from the corresponding aR239:cE111J distance being biased in the eABF calculation. At convergence, uniform sampling is expected in the interval [0.25 nm, 2.5 nm]. For the contact threshold of 0.5 nm used here, this translates into a predicted (0.5 − 0.25)*/*(2.5 − 0.25) ≃ 11% contact probability, which is consistent with the observed values.

**Figure S5:**
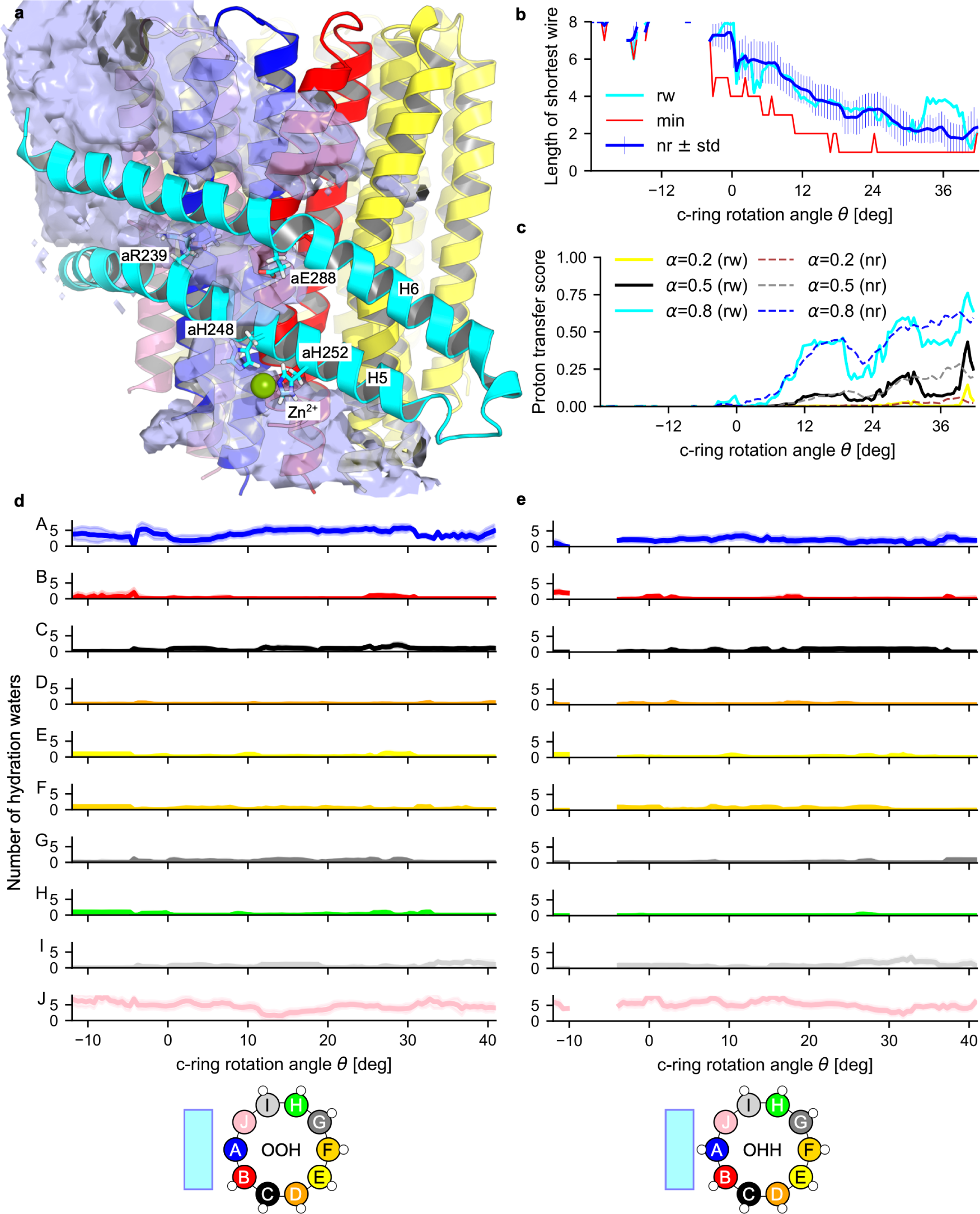
Water-chain analysis. (a) Water populates the a/c interface through the access and exit half-channels. Shown is the water density averaged over the first 45 ns of stratified eABF simulation in state OOH and window *W*_1_, which is representative of the water distribution in the pre-rotation configuration. For clarity, only helices H5 and H6 of the a-subunit are displayed. Water enters the access half-channel through an opening between H5 and H6 in the vicinity of aH248, aH252, the Zn^2+^ ion and aE288. (b) *θ*-dependent length of the shortest water bridge (conditioned on existence). Shown are the eABF-reweighted average (cyan), the minimal sampled value (red) and the non-reweighted average ± standard deviation. (c,d) *θ*-dependent average number of hydration waters for all 10 cE111 side chains in state OOH (c) and OHH (d). Shown are the eABF-reweighted averages ± standard deviation. Sketches at the bottom illustrate protonation states and coloring and labelling conventions. Hydration waters are defined as within 3.4 Å of the carboxylate oxygens.

**Figure S6:**
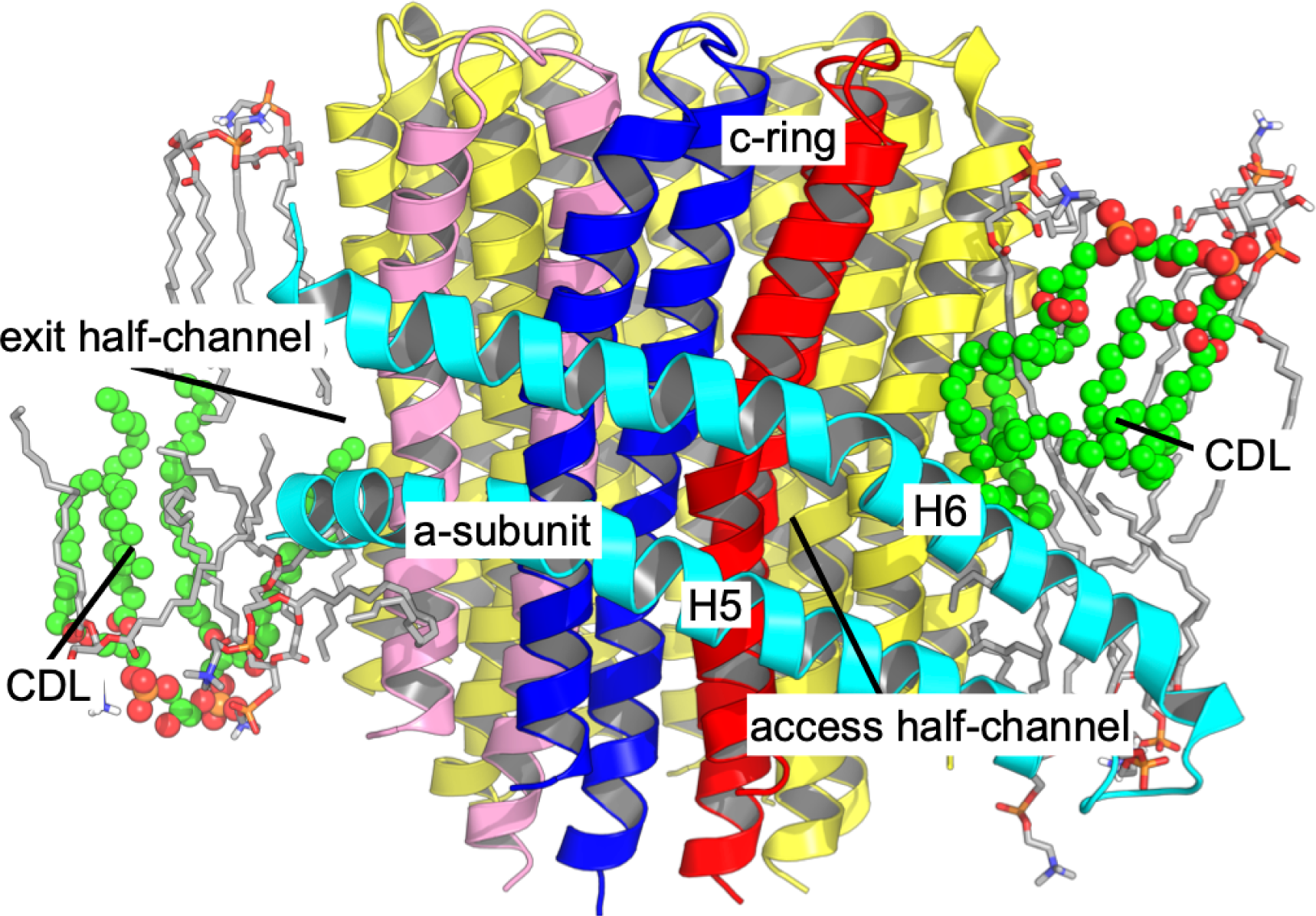
Lipid molecules in the vicinity of the a/c interface. Shown is the equilibrated structure of state OOH as an example of typical lipid distribution. POPE/POPC/POPI (grey sticks) and cardiolipin molecules (CDL, green spheres) mediate a/c contacts.

**Figure S7:**
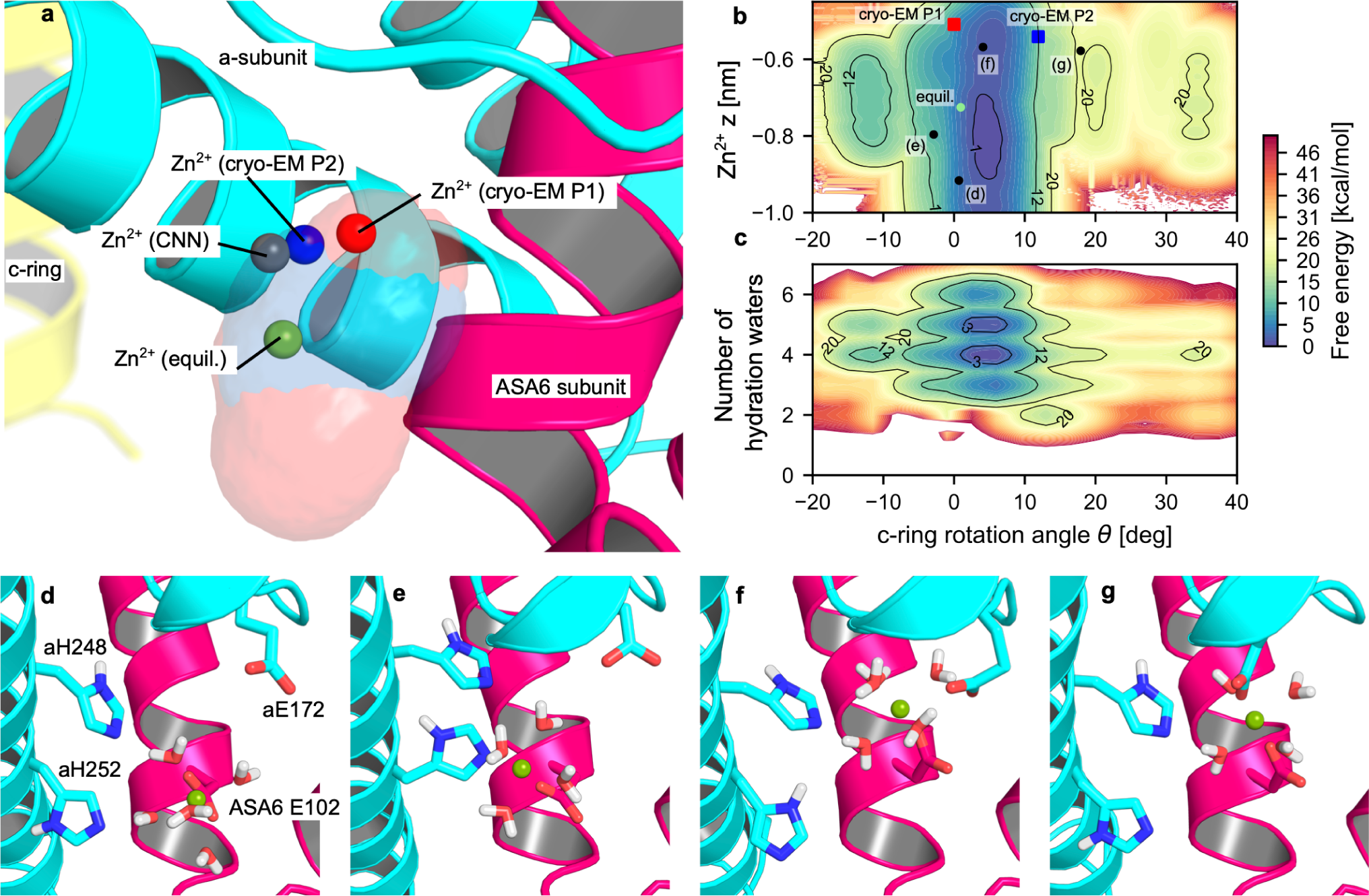
Dynamics of the a-subunit-bound Zn^2+^ ion. (a) Alternate positions explored by Zn^2+^ in cryo-EM and eABF simulations. Shown are the state OOH equilibrated structure, and the Zn^2+^ ion in cryo-EM P1 (red sphere), cryo-EM P2 (blue sphere), convolutional neural network prediction (grey sphere) and state OOH equilibrated structure (green sphere). Also shown are the 0.01 Å^−3^ isocontours of the Zn^2+^ positional probability densities collected by averaging (without reweighting) over the eABF windows *W*_11_ (*i.e.,*, representative of the P1 basin, transparent red surface) and *W*_12_ (*i.e.,*, representative of the P2 basin, transparent blue surface). (b) eABF-reweighted free energy landscape of Zn^2+^ *z*-coordinate (relative to the center of geometry of the ASA6 subunit) versus *θ*. Configurations shown in a. and (d-g) are positioned on the plot. (c). eABF-reweighted free energy landscape of the number of water molecules in the first hydration shell of Zn^2+^ (*i.e.,* within 3 Å) versus *θ*. Gaussian KDE with bandwidth = 0.1 is used to turn discrete hydration numbers into continuous values. Taken together, (b) and (c) show that Zn^2+^ undergoes extensive fluctuations of position (roughly 5 Å of amplitude) and hydration in the OOH ground state. The transition to P2 restricts these fluctuations. (d-g) Typical simulation frames illustrating the alternate positions and hydration patterns of Zn^2+^. Notably, we observe that 1) Zn^2+^ is always interacting directly with residue ASA6 E102, and through a water-mediated interaction with aH248, that 2) downward poses (d-e, *z <* −0.8 nm) entail interactions (either direct or water-mediated) with aH252, but not aE172, whereas 3) the opposite is true for upward poses (f-g, *z >* 0.8 nm). This indicates that the positional dynamics of Zn^2+^ is controlled by competition between aE172 and aH252 for interaction, and that interaction with aE172 is favored when the c-ring is in state P2 through a yet unknown mechanism.

**Figure S8:**
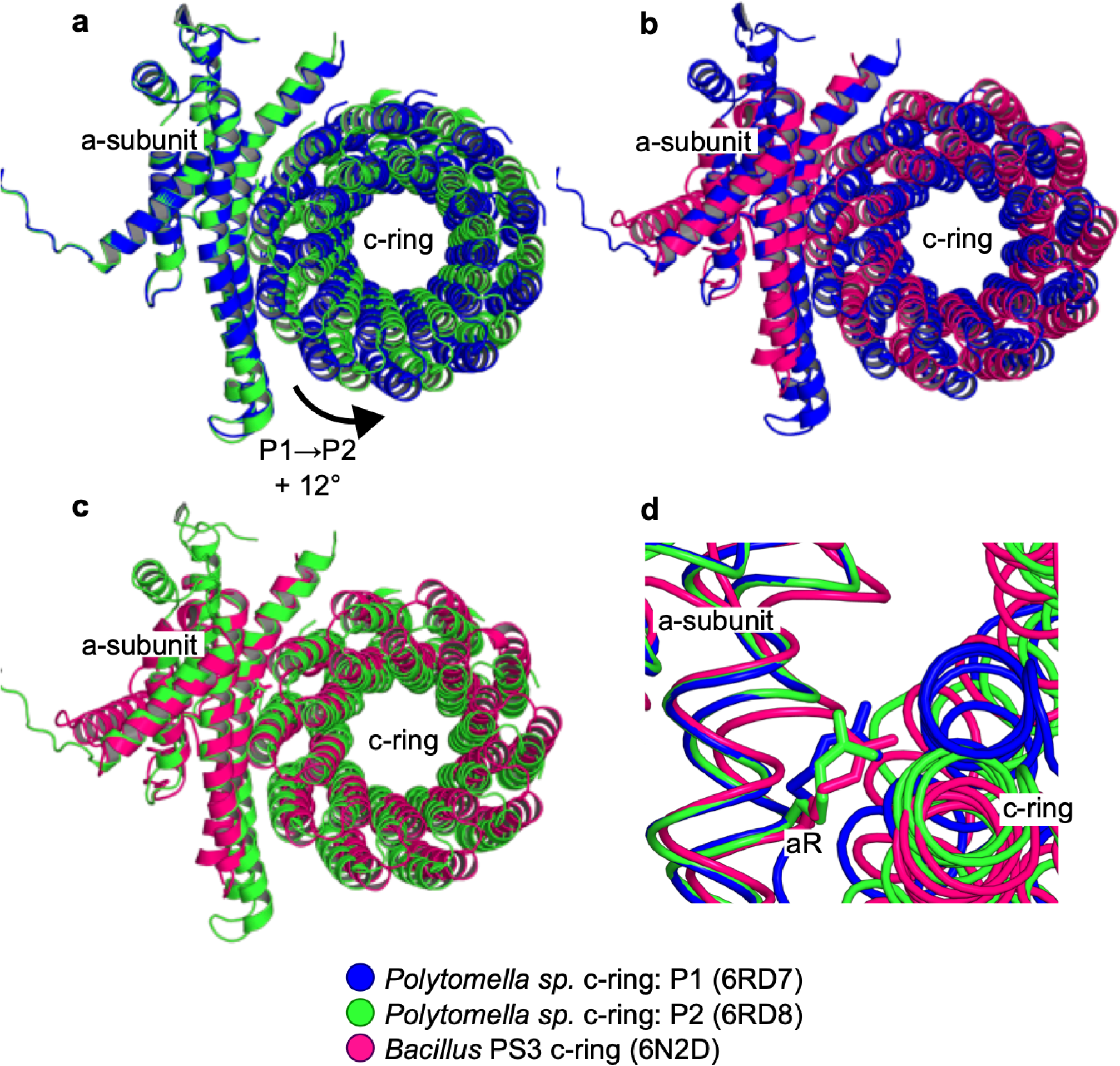
Comparison of c-ring positions in cryo-EM structures of *Polytomella sp.* and *Bacillus sp. PS3* ATP synthases. To isolate c-ring movement, structures are aligned on the structurally conserved a-subunit using Pymol [77] (a) Alternate c-ring positions P1 (6RD7/6RD9, blue) and P2 (6RD8, green) seen in *Polytomella sp.* ATP synthase [33]. P2 is rotated forward by ≈12^◦^ with respect to P1 used as a reference. (b) Comparison of c-ring positions between *Polytomella sp.* P1 structure (6RD7/6RD9, blue) and *Bacillus PS3* structure (6N2D, hotpink) [19] shows that the *Bacillus* c-ring is rotated forward with respect to P1. (c) Comparison of c-ring positions between *Polytomella sp.* P2 structure (6RD8, green) and *Bacillus PS3* structure (6N2D, hotpink) shows that the *Bacillus* c-ring occupies approximately the same position as P2. (d) Close-up on the conserved a-subunit Arginine residue aR (aR239 in *Polytomella sp.*, aR169 in *Bacillus*) showing that the aR CA atoms are superimposed. This illustrates that the structural alignment onto the a-subunit is meaningful and can be used to compare c-ring rotational states across species.

